# Mortality causes universal changes in microbial community composition

**DOI:** 10.1101/396499

**Authors:** Clare I. Abreu, Jonathan Friedman, Vilhelm L. Andersen Woltz, Jeff Gore

## Abstract

All organisms are sensitive to the abiotic environment, and a deteriorating environment can lead to extinction. However, survival in a multispecies community also depends upon inter-species interactions, and some species may even be favored by a harsh environment that impairs competitors. A deteriorating environment can thus cause surprising transitions in community composition. Here, we combine theory and laboratory microcosms to develop a predictive understanding of how simple multispecies communities change under added mortality, a parameter that represents environmental harshness. In order to explain changes in a multispecies microbial system across a mortality gradient, we examine its members’ pairwise interactions. We find that increasing mortality favors the faster grower, confirming a prediction of simple models. Furthermore, if the slower grower outcompetes the faster grower in environments with low or no added mortality, the competitive outcome can reverse as mortality increases. We find that this tradeoff between growth rate and competitive ability is indeed prevalent in our system, allowing for striking pairwise outcome changes that propagate up to multispecies communities. These results argue that a bottom-up approach can provide insight into how communities will change under stress.

## Introduction

Ecological communities are defined by their structure, which includes species composition, diversity, and interactions^1^. All such properties are sensitive to the abiotic environment, which influences both the growth of individual species and the interactions between them. The structure of multispecies communities can thus vary in complex ways across environmental gradients^2–7^. A major challenge is therefore to predict how a changing environment affects competition outcomes and alters community structure. In particular, environmental deterioration can radically change community structure. Instances of such deterioration include antibiotic use on gut microbiota^8^, ocean warming in reef communities^9^, overfishing in marine ecosystems^10^, and habitat loss in human-modified landscapes^11^. Such disturbances can affect community structure in several ways, such as allowing for the spread of invasive species^12^, causing biodiversity loss and mass extinction^13,14^, or altering the interactions between the remaining community members^15,16^. For example, a stable ecosystem can be greatly disrupted by the removal of a single keystone species, potentially affecting species with which it does not directly interact^17–19^.

A common form of environmental deterioration is increased mortality, which can be implemented in the laboratory in a simple way. In fact, the standard method of cultivating and competing bacteria involves periodic dilution into fresh media, a process that necessarily discards cells from the population. The magnitude of the dilution determines the fraction of cells discarded and therefore the added mortality rate, making environmental harshness easy to tune experimentally.

The choice of dilution factor often receives little attention, yet theoretical models predict that an increased mortality rate experienced equally by all species in the community can have dramatic effects on community composition. In particular, it is predicted that such a global mortality will favor the faster-growing species in pairwise competition, potentially reversing competition outcomes from dominance of the slow grower to dominance of the fast grower^1,20,21^. Indeed, there is some experimental support for competitive reversals in chemostat competition experiments between microbial species with different growth rates^22– 24^. A less-explored prediction is that if a high mortality rate causes a competitive reversal, the competition will also result in either coexistence or bistability (where the winner depends on the starting fraction) at some range of intermediate mortality^25–27^. In addition, little is known about how mortality will alter the composition of multispecies communities.

In this paper, we report experimental results that expand upon the prior literature regarding pairwise competition, and we use the pairwise outcomes to develop a predictive understanding of how multispecies community composition changes with increased mortality. First, experimental pairwise competition of five bacterial species confirmed that 1) increased mortality favors the fast grower in a competition, and can reverse the winner of the competition from slow grower to fast grower, and 2) at intermediate dilution rates, either coexistence or bistability occurs. Interestingly, we find that a pervasive tradeoff between growth rate and competitive ability in our system favors slow growers in high-density, low-mortality environments, enabling striking changes in outcomes as mortality increases. Second, to bridge the pairwise results to three- and four-species communities, we employed simple predictive pairwise assembly rules^28^, where we find that the pairwise outcomes such as coexistence and bistability propagate up to the multispecies communities. Our results highlight that the seemingly complicated states a community adopts across a mortality gradient can be traced back to a predictable pattern in the outcomes of its constituent pairs.

## Results

To probe how a changing environment affects community composition, we employed an experimentally tractable system of soil bacteria competitions subject to daily growth/dilution cycles across six dilution factors (Fig. 1A). We selected five species of soil bacteria: *Enterobacter aerogenes* (*Ea*), *Pseudomonas* aurantiaca (*Pa*), *Pseudomonas citronellolis* (*Pci*), *Pseudomonas putida* (*Pp*), and *Pseudomonas veronii* (*Pv*). These species have been used in previous experiments by the group, which did not vary dilution factor^28,29^. All five species grow well in our defined media containing glucose as the primary carbon source (see Methods) and have distinct colony morphology that allows for measuring species abundance by plating and colony-counting on agar.

**Figure 1:**
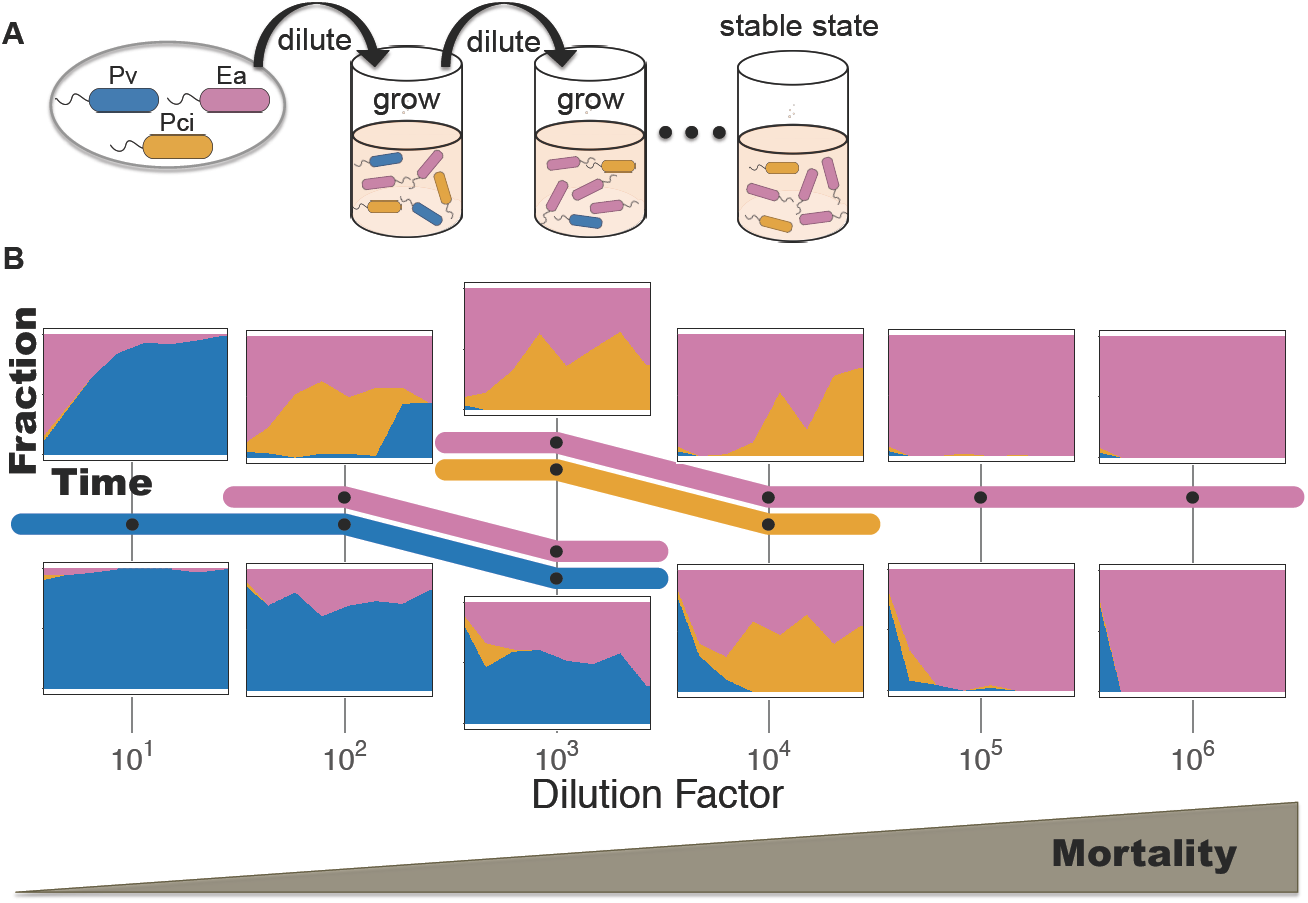
Increasing dilution causes striking shifts in a three-species community. **A)** To probe how added mortality changes community composition, we competed three soil bacteria over a range of dilution rates. Cells were inoculated and allowed to grow for 24 hours before being diluted into fresh media. This process was continued for seven days, until a stable equilibrium was reached. The magnitude of the dilution factor (10 to 10^6^) determines the fraction of cells discarded, and thus the amount of added mortality. **B)** We began with a three-species community (*Enterobacter aerogenes* (*Ea*), *Pseudomonas citronellolis* (*Pci*), and *Pseudomonas veronii* (*Pv*)), initialized from four starting fractions at each dilution factor. The outcomes of two of the starting fractions are shown, along with a “subway” map, where survival of species is represented with colors assigned to each species. Black dots indicate where data was collected, while colors indicate the range over which a given species is inferred to survive. Species *Pv* dominates at the lowest dilution factor, and *Ea* dominates at the highest dilution factors. The stacking of two colors represents coexistence of two species, whereas the two levels at dilution factor 10^3^ indicate bistability, where both coexisting states, *Ea*-*Pv* and *Ea*-*Pci*, are stable and the starting fraction determines which stable state the community reaches.

We began by competing three of the five species, *Ea, Pci*, and *Pv*, for seven 24-hour cycles under six different dilution factor regimes. To assay for alternative stable states, each dilution factor condition was initialized by four different starting fractions (equal abundance as well as prevalence of one species in a 90-5-5% split). Despite the simplicity of the community and the experimental perturbation, we observed five qualitatively different outcomes corresponding to different combinations of the species surviving at equilibrium (Fig. 1B). At the highest and lowest dilution factors, one species excludes the others at all starting fractions (*Pv* at low dilution, *Ea* at high dilution). Two coexisting states (*Ea*-*Pv* and *Ea*-*Pci*) occur at medium low (10^2^) and medium high (10^4^) dilution factors, again independent of the starting fractions of the species. However, at intermediate dilution factor (10^3^), we found that the surviving species depended upon the initial abundances of the species. At this experimental condition, the system displays bistability between the two different coexisting states (*Ea*-*Pv* and *Ea*-*Pci*) that were present at neighboring dilution factors. These three species therefore display a surprisingly wide range of community compositions as the mortality rate is varied.

To make sense of these transitions in community composition, we decided to first focus on two-species competitions, not only because they should be simpler, but also because prior work from our group gives reason to believe that pairwise outcomes are sufficient for predicting multispecies states^28^. Accordingly, we used a simple two-species Lotka-Volterra competition model with an added mortality term *δN*_*i*_ experienced equally by both species^21^:

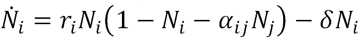

where *N*_*i*_ is the density of species *i* (normalized to its carrying capacity), *r*_*i*_ is the maximum growth rate of species *i*, and the competition coefficient *α*_*ij*_ is a dimensionless constant reflecting how strongly species *i* is inhibited by species *j* (Fig. 2). This model can be re-parameterized into the Lotka-Volterra model with no added mortality, where the new competition coefficients 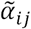 now depend upon *r*_*i*_ and *δ* (see S2 for derivation):

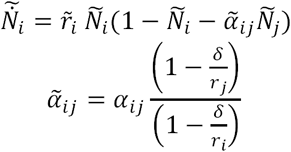

**Figure 2:**
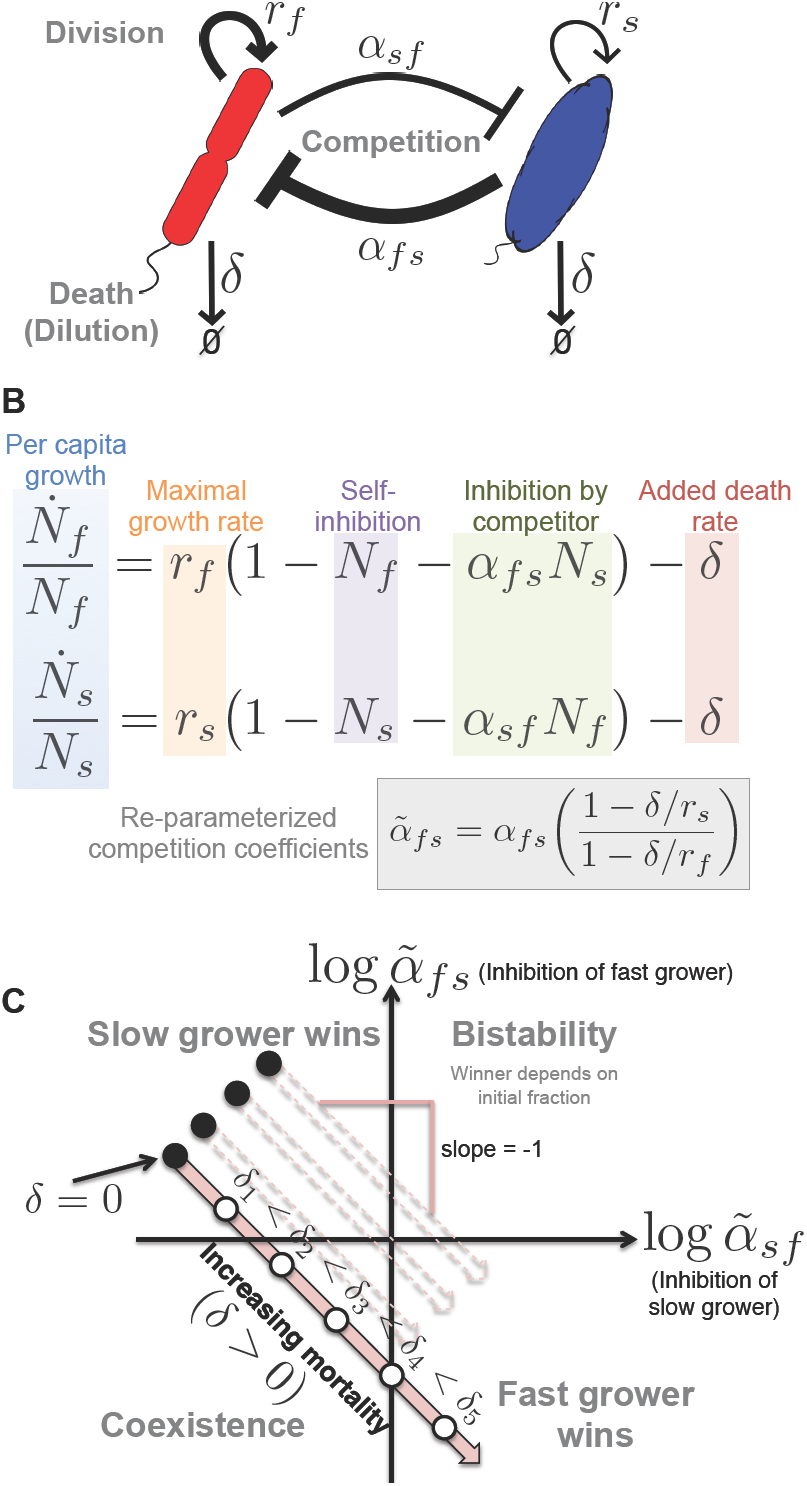
An increasing global mortality rate is predicted to favor the fast grower. **A-B)** Here we illustrate the parameters of the Lotka-Volterra (LV) interspecific competition model with added mortality: population density *N*, growth *r*, death *δ* (subscript *f* for fast grower and *s* for slow grower), and the strengths of inhibition *α*_*sf*_ and *α*_*fs*_. The width of arrows in (**A**) corresponds to an interesting case that we observe experimentally, in which the fast grower is a relatively weak competitor. **C)** The outcomes of the LV model without mortality depend solely upon the competition coefficients *α*, and the phase space is divided into one quadrant per outcome. If it is a strong competitor, the slow grower can exclude the fast grower. Imposing a uniform mortality rate *δ* on the system, however, favors the faster grower by making the re-parameterized competition coefficients 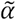 depend on *r* and *δ*. Given that a slow grower dominates at low or no added death, the model predicts that coexistence or bistability will occur at intermediate added death rates before the outcome transitions to dominance of the fast grower at high added death (see S2 for derivation).

The outcome of competition—dominance, coexistence, or bistability—simply depends upon whether each of the 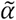 are greater or less than one, as in the basic Lotka-Volterra competition model^21^.

In this model, it is possible for a slow grower (*N*_*s*_) to outcompete a fast grower (*N*_*f*_) if the slow grower is a strong competitor (*α*_*fs*_ *f* 1) and the fast grower is a weak competitor (*α*_*sf*_ < 1) (Fig. 2). However, the competition coefficients change with increasing mortality *δ* in a way that favors the fast grower: 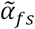 shrinks and 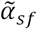 grows, eventually leading the fast grower to outcompete the slow grower. A powerful way to visualize this change is to plot the outcomes, as determined by the competition coefficients (Fig 2C); increasing mortality causes the outcome to traverse a 45° trajectory through the phase space, leading to the fast grower winning at high mortality. At intermediate mortality, the model predicts the two species will either coexist or be bistable. This model therefore makes very clear predictions regarding how pairwise competition will change under increased mortality, given the aforementioned slow grower advantage at low mortality.

To test these predictions in the laboratory, we performed all pairwise competitions at multiple dilution factors and starting fractions of our five bacterial species: *Pp, Ea, Pci, Pa, Pv* (listed in order from fast to slow growing species). We find that these pairwise competitive outcomes change as expected from the LV model, where increased dilution favors the fast grower (Fig S1). For example, in *Ea*-*Pv* competition we find that *Pv*, despite being the slower grower, is able to exclude *Ea* at low dilution rates (Fig 3B, left panel). From the standpoint of the LV model, *Pv* is a strong competitor despite being a slow grower in this environment. However, as predicted by the model, at high dilution rates the slow-growing *Pv* is excluded by the fast-growing *Ea* (Fig 3B, right panel). Importantly, *Pv* is competitively excluded at a dilution factor of 10^4^, an experimental condition at which it could have survived in the absence of a competitor. Finally, and again consistent with the model, at intermediate dilution rates we find that the *Ea*-*Pv* pair crosses a region of coexistence, where the two species reach a stable fraction over time that is not a function of the starting fraction (Fig 3B, middle panel). The *Ea*-*Pv* pair therefore displays the transitions through the LV phase space in the order predicted by our model (Fig 3A-D).

**Figure 3:**
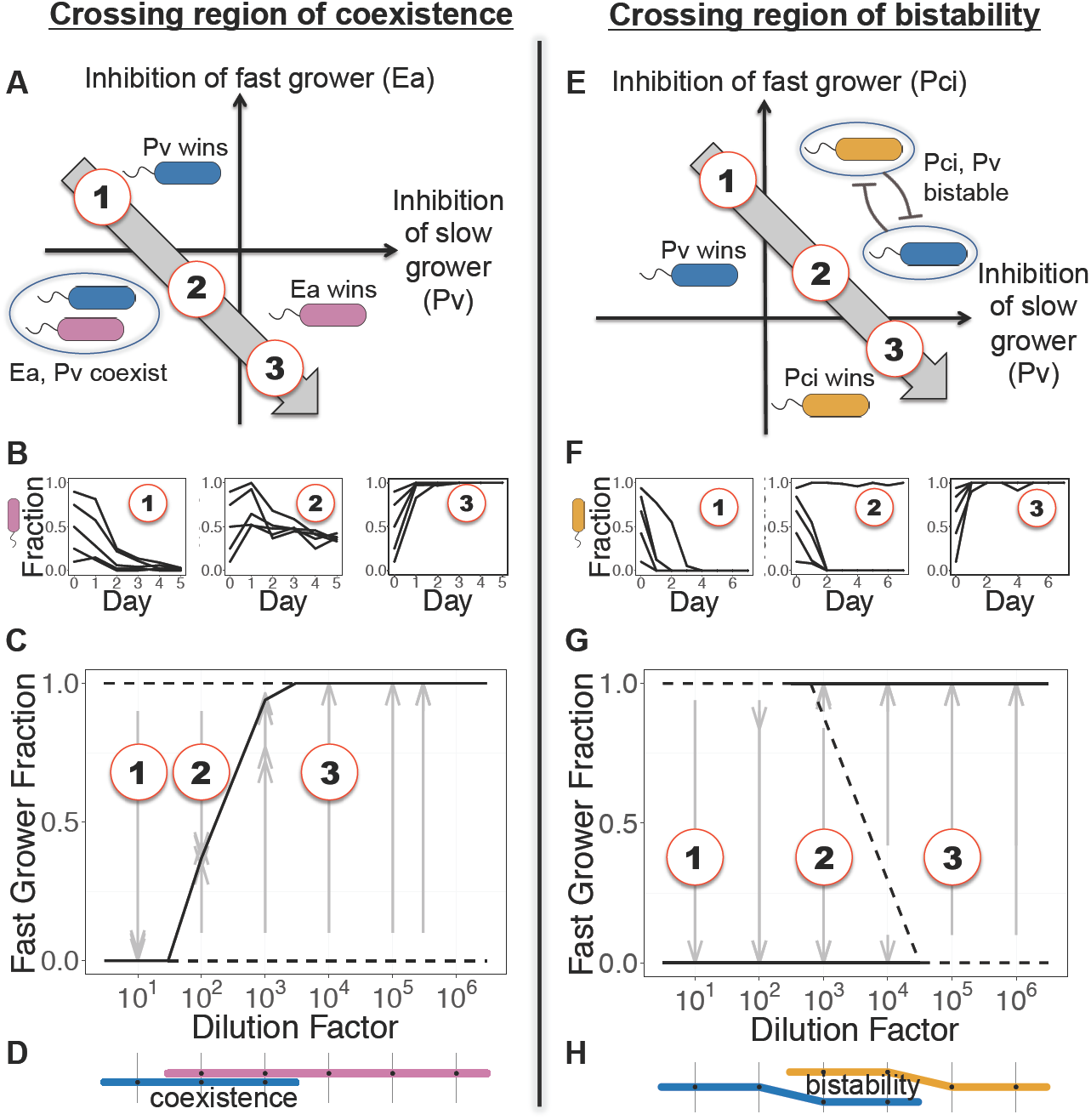
In pairwise competition experiments, increasing dilution favors the faster grower, and coexistence or bistability occur at intermediate dilution. **A)** Experimental results are shown from a competition between *Pv* (blue) and *Ea* (pink). **B)** <u>Left</u> <u>pane</u>l: Despite its slow growth rate, *Pv* excludes faster grower *Ea* at the lowest dilution factor. <u>Middle pane</u>l: Increasing death rate causes the outcomes to traverse the coexistence region of the phase space. <u>Right pane</u>l: As predicted, fast-growing *Ea* dominates at high dilution factor. **C)** An experimental bifurcation diagram shows stable points with a solid line, and unstable points with a dashed line. The stable fraction of coexistence shifts in favor of the fast grower as dilution increases. Gray arrows show experimentally measured time trajectories, beginning at the starting fraction and ending at the final fraction. **D)** A “subway map” denotes survival/extinction of a species at a particular dilution factor with presence/absence of the species color. **E-F)** *Pv* outcompeted another fast grower *Pci* (yellow) at low dilution factors, but the pair became bistable instead of coexisting as dilution increased; the unstable fraction can be seen to shift in favor of the fast grower **(G)**. **H)** Two levels in the subway map show bistability.

The LV model predicts that other pairs will cross a region of bistability rather than coexistence, and indeed this is what we observe experimentally with the *Pci*-*Pv* pair (Fig 3E-H). Once again, the slow-growing *Pv* dominates at low dilution factor yet is excluded at high dilution factor. However, at intermediate dilution factors this pair crosses a region of bistability, in which the final outcome depends upon the starting fractions of the species. The Lotka-Volterra model with added mortality therefore provides powerful insight into how real microbial species compete, despite the many complexities of the growth and interaction that are necessarily neglected in a simple phenomenological model.

Indeed, a closer examination of the trajectory through the LV phase space of the *Pci*-*Pv* pair reveals a violation of the simple outcomes allowed within the LV model. In particular, at dilution factor 10^2^ we find that when competition is initiated from high initial fractions of *Pci* that *Pv* persists at low fraction over time (Fig. 3G). This outcome, a bistability of coexistence and exclusion (rather than of exclusion and exclusion), is not an allowed outcome within the LV model (modifications to the LV model can give rise to it, as shown by ^30^). This subtlety highlights that the transitions (e.g. bifurcation diagrams in Fig 3C,G) can be more complex than what occurs in the LV model, but that nonetheless the transitions within the LV model represent a baseline to which quantitative experiments can be compared.

The model predicts that mortality will reverse competition outcomes if and only if a slow grower outcompetes a fast grower at low or no added death, exhibiting a tradeoff between growth and competitive ability. Changes in outcome are therefore most dramatic when a strongly competing slow grower causes the trajectory to begin in the upper left quadrant of the phase space (Fig. 3A, E), allowing it to move through other quadrants as mortality increases. Indeed, in the pairwise competitions described above, the slowest-growing species, *Pv*, is a strong competitor at low dilution factor. To probe this potential tradeoff more extensively, we compared the growth rates of our five species in monoculture (Fig. S3) to their competitive performance at low dilution factor. In seven of the ten pairs, the slower grower excluded the faster grower, and the other three pairs coexisted (Fig. S1). We therefore find that our five species display a pervasive tradeoff between growth rate and competitive ability, possibly because the slower-growing species fare better in high-density environments that reach saturation.

To visualize how competitive success changes with dilution factor, we defined the competitive score of each species to be its mean fraction after reaching equilibrium in all pairs in which it competed. The aforementioned tradeoff can be seen as an inverse relationship between growth rate and competitive score at the lowest dilution factor (Fig. 4A). As predicted, the performance of the fast-growing species increases monotonically with increasing dilution factors (Fig. 4B). Competitive superiority of the slowest grower (*Pv*) at low dilution rates transitions to the next-slowest (*Pa*) at intermediate rates, before giving rise to dominance of the fastest growers (*Pci, Ea, Pp*) at maximum rates (Fig 4B-D). We therefore find that the mortality rate largely determines the importance of a species’ growth rate to competitive performance in pairwise competition.

**Figure 4:**
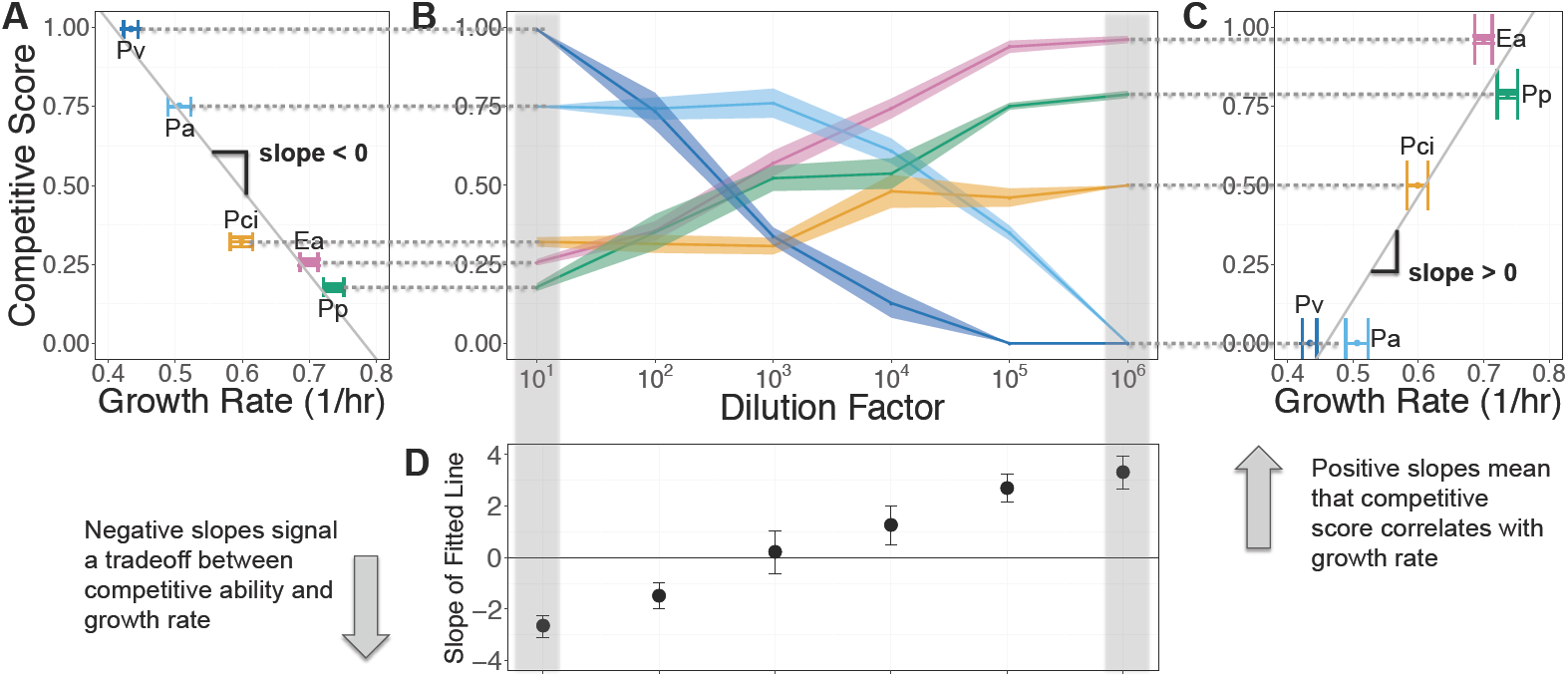
In experimental competition of five bacterial species in pairs, a tradeoff between growth and competitive ability leads to strong dependence of outcome on dilution factor. The LV model predicts that increasing dilution will favor faster-growing species over slower-growing ones. If fast growers dominate at low dilution actors, though, no changes in outcome will be expected. Changes in outcome are therefore most dramatic when slow growers are strong competitors at low dilution, exhibiting a tradeoff between growth rate and competitive ability. **A)** This tradeoff was pervasive in our system: slower growth rates resulted in higher competitive scores at the lowest dilution factor. Growth rate was calculated with OD600 measurements of the time taken to reach a threshold density within the exponential phase; error bars represent the SEM of replicates (n ∼ 20 replicates) (Fig. S3). Competitive score was calculated by averaging fraction of a given species across all pairwise competitive outcomes; error bars were calculated by bootstrapping, where replicates of mean experimental outcomes of a given pair were sampled with replacement. **B)** The competitive scores in **A** are extended to all dilution factors. The slowest grower’s score monotonically decreases with dilution, while the fast growers’ scores increase, and an intermediate grower peaks at intermediate dilution factor. **C)** At high dilution factors, the order of scores is reversed. **D)** At low dilution factors 10 and 10^2^, competitive ability is negatively correlated with growth rate; the correlation becomes positive above dilution factor 10^3^. Error bars are the standard error coefficients given by the linear regression function lm in R.

Now that we have an understanding of how pairwise competitive outcomes shift in response to increased mortality, we return to the seemingly complicated set of outcomes observed in our original three-species community (Fig. 1). In a previous study^28^, we developed community assembly rules that allow for prediction of species survival in multispecies competition from the corresponding pairwise competition outcomes. These rules state that in a multispecies competition, a species will survive if and only if it coexists with all other surviving species in pairwise competition. If one or more bistable pairs is involved in a multispecies community, the assembly rules allow for either of the stable states. We see that the seemingly complicated trio outcomes follow from these assembly rules applied to our corresponding pairwise outcomes at all dilution factors (Fig. 5). For example, at the lowest dilution factor (10), *Ea*-*Pci* coexist, but each of these species is excluded by *Pv* in pairwise competition, thus leading to the (accurate) prediction that only *Pv* will survive in the trio competition. In addition, we observe that the bistability of *Pci*-*Pv* at dilution factor 10^3^ propagates up to lead to bistability in the trio, but with each stable state corresponding to coexistence of two species. The only trio outcome not successfully predicted by the rules is the occasional persistence of *Pci* at a dilution factor of 10^5^ (Figs. 5D, S8). Our analysis of pairwise shifts under increased mortality therefore provides a predictive understanding of the complex shifts observed within a simple three-species bacterial community.

**Figure 5:**
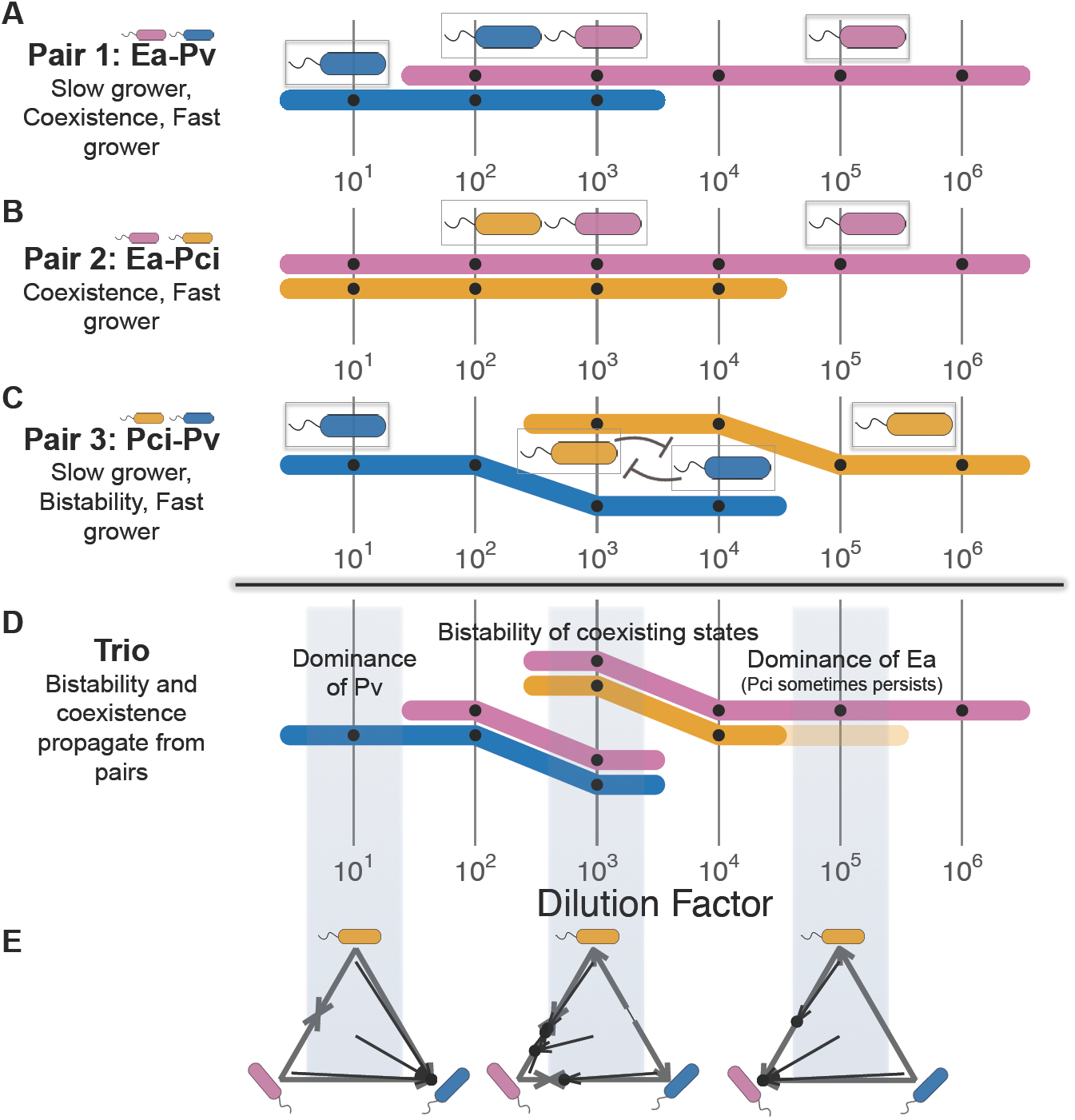
Coexistence and bistability propagate from pair to trio, as predicted by assembly rules. **A-C)** “Subway maps” show pairwise competition outcome trajectories across changing dilution factor, as explained in Figs. 1 and 3. The fast grower’s line is always plotted above the slow grower’s line. Of the three pairs that make up the community *Ea*-*Pci*-*Pv*, two are coexisting (**A**, **B**) and one is bistable (**C**). **D)** The pairwise assembly rules state that a species will survive in a community if it survives in all corresponding pairs. At dilution factor 10, *Ea* and *Pci* coexist, but both are excluded by *Pv*. The rules correctly predict that *Pv* will dominate in the trio. Because both species can be excluded in a bistable pair, a bistable pairwise outcome propagates to the trio as more than one allowed state. Each of the bistable species can be seen separately coexisting with *Ea* at dilution factor 10^3^, as they do in pairs. The assembly rules failed at 10^5^ for one out of four starting conditions: *Pci* coexists with *Ea* when it should go extinct (Fig. S8). **E)** Three-species competition results are shown in simplex plots. Arrows begin and end at initial and final fractions, respectively. Edges represent pairwise results, and black dots represent trio results.

To determine whether our analysis of community shifts under mortality is more broadly applicable, we combined our five species into various three- and four-species subsets, similar to the *Ea*-*Pci*-*Pv* competition (Fig. 5). In total, we competed five three-species communities and three four-species communities at all six dilution factors. Overall, a quantitative generalization of our assembly rules (see Methods) predicted the equilibrium fractions with an error of 14%, significantly better than the 39% error that results from predictions obtained from monoculture carrying capacity (Table 1, Fig. S2). These results indicate that pairwise outcomes are good predictors of multispecies states in the presence of increased mortality.

**Table 1:**
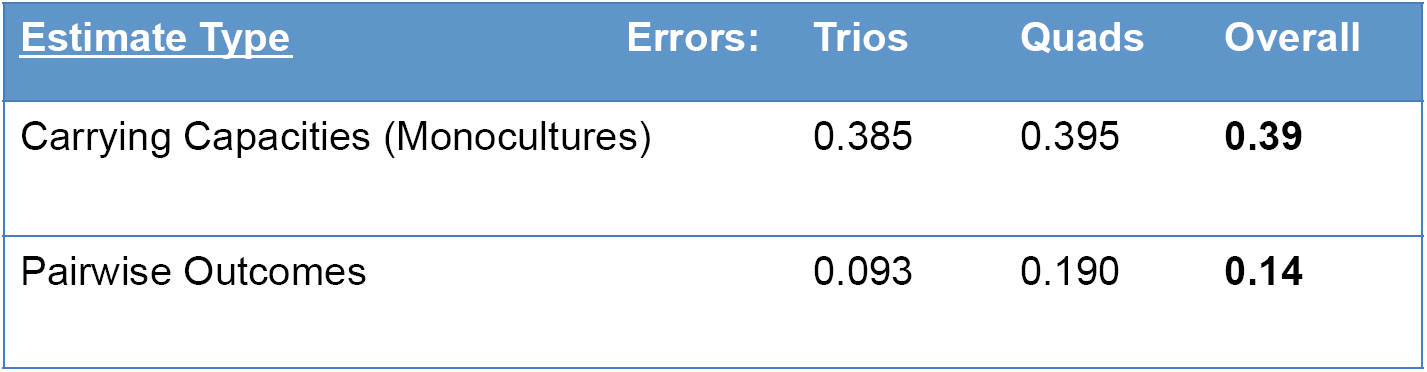
Errors of pairwise assembly rules are much lower than errors of estimates using monoculture carrying capacity. We made quantitative predictions of the relative fractions in multispecies competition outcomes using both monoculture carrying capacities as well as the pairwise assembly rules. Errors of quantitative predictions are the L2 norm of the distance between predicted fixed point and observed fixed point (see Methods and Fig. S2). The values shown are mean error normalized by the maximum error.

## Discussion

The question of how community composition will change in a deteriorating environment is essential, as climate change, ocean acidification, and deforestation infringe upon many organisms’ habitats, increasing mortality either directly, by decimating populations, or indirectly, by making the environment less hospitable to them. We used an experimentally tractable microbial microcosm to tune mortality through dilution rate, and found a pervasive tradeoff between growth rate and competitive ability (Fig. 4). This tradeoff causes slow growers to outcompete fast growers in high-density, low-dilution environments. Increasing mortality favors fast growers, in line with model predictions. We observed coexistence and bistability at intermediate dilution factors in pairwise experiments (Fig. 3), and found that such coexistence and bistability propagated up to three- and four-species communities (Fig. 5). Coexistence was more common than bistability, which is in line with expectations of optimal foraging theory^1^. We were able to explain seemingly complicated multispecies states with pairwise results, which traversed all possible competition outcomes allowed by the two-species model.

The aforementioned tradeoff made for striking transitions in the communities that we studied. Without the tradeoff, the model would be less useful. If a fast grower outcompetes a slow grower at low dilution rates, the model predicts no change in outcome at higher dilution rates. Our results at low dilution are consistent with previous experimental evidence of a tradeoff between growth and competitive ability among different mutants of the same bacterial strain^31^ and protozoa^32^. Additionally, data show that antibiotic resistance, despite its clear competitive benefit, imposes a fitness cost on bacteria^33^. There is also evidence that seed-producing plants exhibit a growth/competition tradeoff; plants that produce larger seeds necessarily produce fewer of them, but were found to have better seedling establishment when competing with smaller-seeded plants^34,35^.

The mechanism for the competitive ability of the slow growers in our system is not easily explained; supernatant experiments, in which fast growers were placed in slow growers’ filtered spent media showed little or no inhibition of growth compared to controls (Fig. S6). Furthermore, in monocultures the slow growers exhibited higher lag times than the fast growers (Fig. S5), which would seem to be disadvantageous in low-dilution, high-density conditions where resources could be quickly consumed by a competitor with a shorter lag^36^. The frequency of the tradeoff in other systems is a question worthy of further investigation, in particular because natural microbial systems, such as soil communities or the gut microbiome, are better represented with a low dilution rate than a high dilution rate^37,38^.

We employed a set of simple pairwise assembly rules^28^ to predict the states of three- and four-species communities (Table 1, Fig. S2). The rules’ success is in line with recent microbial experiments suggesting that pairwise interactions play a key role in determining multispecies community assembly^28,39^ and community-level metabolic rates^40^; in contrast, some theory and empirical evidence supports the notion of pervasive and strong higher-order interactions^41– 44^. Our results provide support for a bottom-up approach to simple multispecies communities, and show that pairwise interactions alone can generate multispecies states that appear nontrivial. In the model, this happens because up to three qualitative regimes of pairwise competition translate to more possible combinatorial multispecies outcomes.

Here we found that the LV model with added mortality provided useful guidance for how experimental competition would shift under increased dilution, but resource-explicit models may in some cases provide additional mechanistic insight^45,46^. In particular, various resource-explicit models can recapitulate the qualitative changes predicted by the LV model with added mortality. For example, the R* rule states the species that can survive on the lowest equilibrium resource concentration will dominate other species^1^. The equilibrium concentration increases with the dilution rate, thus favoring the species with the highest maximal growth rate (see S4). However, a species with a low maximal rate may dominate under low dilution if it can grow more efficiently at low resource concentrations. Resource explicit models are most commonly used for simple environments, whereas here we worked with media containing three carbon sources. In addition, we have found that complex media with many dozens of carbon sources yields similar changes in pairwise outcomes with dilution and multispecies communities that could be predicted by the pairwise assembly rules (Figs. S2,7). Further work is necessary to explore the circumstances in which phenomenological or resource-explicit models should be used^47–49^.

It is also important to note that not all deteriorating environments will cause such simple and uniform increases in mortality. Antibiotics, and in particular *β*-lactam antibiotics, might selectively attack fast growers over slow growers^50^. Overfishing might target certain species of fish. Climate change might affect growth rate rather than death rate by increasing temperature, which usually increases growth rates^51^. In such a case, it is not certain whether environmental deterioration in the form of warming would favor slow growers or fast growers. An important direction for future research is to determine whether other changes to the environment will have similarly simple consequences for the composition of microbial communities. In this study, we have seen how a simple prediction about a simple perturbation in pairwise competition—increased mortality will favor the faster-growing species—allowed us to interpret seemingly nontrivial outcomes in multispecies communities.

## Methods

### Species and media

The soil bacterial species used in this study were *Enterobacter aerogenes* (Ea, ATCC#13048), *Pseudomonas aurantiaca* (Pa, ATCC#33663), *Pseudomonas citronellolis* (Pci, ATCC#13674), *Pseudomonas putida* (Pp, ATCC#12633) and *Pseudomonas veronii* (Pv, ATCC#700474). All species were obtained from ATCC. Two types of growth media were used: one was complex and undefined, while the other was minimal and defined. All results presented in the main text are from the defined media. All species grew in monoculture in both media. The complex medium was 0.1X LB broth (diluted in water). The minimal medium was S medium, supplemented with glucose and ammonium chloride. It contains 100 mM sodium chloride, 5.7 mM dipotassium phosphate, 44.1 mM monopotassium phosphate, 5 mg/L cholesterol, 10 mM potassium citrate pH 6 (1 mM citric acid monohydrate, 10 mM tri-potassium citrate monohydrate), 3 mM calcium chloride, 3 mM magnesium sulfate, and trace metals solution (0.05 mM disodium EDTA, 0.02 mM iron sulfate heptahydrate, 0.01 mM manganese chloride tetrahydrate, 0.01 mM zinc sulfate heptahydrate, 0.01 mM copper sulfate pentahydrate), 0.93 mM ammonium chloride, 10 mM glucose. 1X LB broth was used for initial inoculation of colonies. For competitions involving more than two species, plating was done on 10 cm circular Petri dishes containing 25 ml of nutrient agar (nutrient broth (0.3% yeast extract, 0.5% peptone) with 1.5% agar added). For pairwise competitions, plating was done on rectangular Petri dishes containing 45 ml of nutrient agar, onto which diluted 96-well plates were pipetted at 10 ul per well.

### Growth rate measurements

Growth curves were captured by measuring the optical density of monocultures (OD 600 nm) in 15-minute intervals over a period of ∼50 hours (Fig. S3). Before these measurements, species were grown in 1X LB broth overnight, and then transferred to the experimental medium for 24 hours. The OD of all species was then equalized. The resulting cultures were diluted into fresh medium at factors of 10^-8^ to 10^-3^ of the equalized OD. Growth rates were measured by assuming exponential growth to a threshold of OD 0.1, and averaging across many starting densities and replicates (n = 19 for *Pci*, n = 22 for all other species). This time-to-threshold measurement implicitly incorporates lag times, because a species with a time lag will take longer to reach the threshold OD than another species with the same exponential rate but no lag time. We also estimated lag times and exponential rates explicitly (Fig. S4). We used these measurements to develop an alternative to the time-to-threshold rates, which also incorporated lag time. To estimate this effective growth rate, we multiplied the exponential rate by a factor depending on lag time and time between daily dilutions (Fig. S5B, S4). This method does change growth rate estimates slightly, but does not change the order of growth rates among the five species, and thus the qualitative predictions of the model (Fig. S5A-B). For this reason, we preferred to use the time-to-threshold method, because it involved only one measurement, rather than two, and had a lower error.

### Competition experiments

Frozen stocks of individual species were streaked out on nutrient agar Petri dishes, grown at room temperature for 48 h and then stored at 4 °C for up to two weeks. Before competition experiments, single colonies were picked and each species was grown separately in 50 ml Falcon tubes, first in 5 ml LB broth for 24 h and next in 5 ml of the experimental media for 24 h. During the competition experiments, cultures were grown in 500 μl 96-well plates (BD Biosciences), with each well containing a 200-μl culture. Plates were incubated at 25°C and shaken at 400 rpm, and were covered with an AeraSeal film (Sigma-Aldrich). For each growth–dilution cycle, the cultures were incubated for 24 h and then serially diluted into fresh growth media. Initial cultures were prepared by equalizing OD to the lowest density measured among competing species, mixing by volume to the desired species composition, and then diluting mixtures by the factor to which they would be diluted daily (except for dilution factor 10^-6^, which began at 10^-5^ on Day 0, to avoid causing stochastic extinction of any species). Relative abundances were measured by plating on nutrient agar plates. Each culture was diluted in phosphate-buffered saline prior to plating. For competitions involving more than two species, plating was done on 10 cm circular Petri dishes. For pairwise competitions, plating was done on 96-well-plate-sized rectangular Petri dishes containing 45 ml of nutrient agar, onto which diluted 96-well plates were pipetted at 10 ul per well. Multiple replicates of the latter dishes were used to ensure that enough colonies could be counted. Colonies were counted after 48 h incubation at room temperature. The mean number of colonies counted, per plating, per experimental condition, was 42.

### Assembly rule predictions and accuracy

In order to make predictions about three- and four-species states, we used the qualitative and quantitative outcomes of pairwise competition. The two types of pairwise outcomes allowed for two types of predictions. First, the qualitative outcomes (dominance/exclusion, coexistence, or bistability) of the pairs were used to predict whether a species would be present or absent from a community. These outcomes are shown in the “subway maps” of Fig. S1, where the presence of a species is noted by the presence of its assigned color. Coexistence is shown by two stacked colors, and bistability is shown by two separated colors. The qualitative error rate is the percentage of species, out of the total number of species (three for trios, four for quads), that are incorrectly predicted to be present or absent (Table 1, Fig. S2-A,B). The qualitative success rate is the percentage of species that are correctly predicted as present or absent (Fig. S2-D).

Second, the quantitative outcomes of the pairs were used to predict the quantitative outcomes of three- and four-species communities. These outcomes are shown in the fraction plots of Fig. S1, where equilibrium points are indicated by the black dots. When two or more species coexist in pairs, the assembly rules predicts they will coexist in multispecies communities, provided that an additional species does not exclude them. The predicted equilibrium coexisting fraction of two species is the same in a community as it is in a pair, while the fractions of more than two coexisting species are predicted with the weighted geometric mean of pairwise coexisting fractions. For example, in a three-species coexisting community, the fraction of species 1 depends on its coexisting fractions with the other two species in pairs:

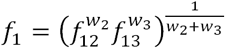

where *f*_12_ is the fraction of species 1 after reaching equilibrium in competition with species 2, 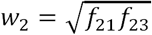 and 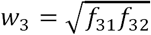. Finally, these predictions are normalized by setting 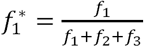.The quantitative error of a particular community outcome is the distance of the predicted fractions from the observed community fractions, measured with the L2 norm. The maximum error, for any number of species, is 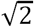, which occurs when a species that was predicted to go extinct in fact dominates:

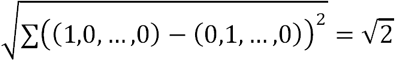

To calculate the overall quantitative errors (Table 1, Fig. S2-C), we divided each error by 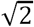 and took the mean.

Finally, we also predicted multispecies states using carrying capacities as measured in monocultures. We assumed that in competition, each species would grow to a density proportionate to its carrying capacity. In other words, the monoculture prediction assumes that all species always coexist. The error from the prediction to the observed data was calculated with the L2 norm, as above.

## Code availability

The code used for analyzing data is available from the first author upon request.

## Data availability

Access to the data is publicly available at TBD.

## References

1. Tilman, D. Resource Competition and Community Structure. (Princeton University Press, 1982).

2. Wellborn, G. A., Skelly, D.K. & Werner, E. E. Mechanisms Creating Community Structure Across a Freshwater Habitat Gradient. Annu. Rev. Ecol. Syst. 27, 337–363 (1996).

3. DÍez, I., Secilla, A., Santolaria, A. & Gorostiaga, J. M. Phytobenthic Intertidal Community Structure Along an Environmental Pollution Gradient. Mar. Pollut. Bull. 38, 463–472 (1999).

4. Yergeau, E. et al. Size and structure of bacterial, fungal and nematode communities along an Antarctic environmental gradient. FEMS Microbiol. Ecol. 59, 436–451 (2007).

5. Lessard, J.-P., Sackett, T. E., Reynolds, W. N., Fowler, D.A. & Sanders, N.J. Determinants of the detrital arthropod community structure: the effects of temperature and resources along an environmental gradient. Oikos 120, 333–343 (2011).

6. Cornwell, W.K. & Ackerly, D.D. Community assembly and shifts in plant trait distributions across an environmental gradient in coastal California. Ecol. Monogr. 79, 109–126 (2009).

7. Mykrä, H., Tolkkinen, M. & Heino, J. Environmental degradation results in contrasting changes in the assembly processes of stream bacterial and fungal communities. Oikos 126, 1291–1298 (2017).

8. Dethlefsen, L. & Relman, D.A. Incomplete recovery and individualized responses of the human distal gut microbiota to repeated antibiotic perturbation. Proc. Natl. Acad. Sci. 108, 4554–4561 (2011).

9. Wernberg, T. et al. Climate-driven regime shift of a temperate marine ecosystem. Science 353, 169–172 (2016).

10. Daskalov, G. M., Grishin, A. N., Rodionov, S. & Mihneva, V. Trophic cascades triggered by overfishing reveal possible mechanisms of ecosystem regime shifts. Proc. Natl. Acad. Sci. 104, 10518–10523 (2007).

11. Larsen, T. H., Williams, N.M. & Kremen, C. Extinction order and altered community structure rapidly disrupt ecosystem functioning. Ecol. Lett. 8, 538–547 (2005).

12. Iii, F. S. C. et al. Consequences of changing biodiversity. Nature (2000). doi:10.1038/35012241

13. Thomas, C. D. et al. Extinction risk from climate change. Nature 427, 145–148 (2004).

14. Bálint, M. et al. Cryptic biodiversity loss linked to global climate change. Nat. Clim. Change 1, 313–318 (2011).

15. Harrington, R., Woiwod, I. & Sparks, T. Climate change and trophic interactions. Trends Ecol. Evol. 14, 146–150 (1999).

16. Ockendon, N. et al. Mechanisms underpinning climatic impacts on natural populations: altered species interactions are more important than direct effects. Glob. Change Biol. 20, 2221–2229 (2014).

17. Paine, R. T. The Pisaster-Tegula Interaction: Prey Patches, Predator Food Preference, and Intertidal Community Structure. Ecology 50, 950–961 (1969).

18. Bond, W. J. Keystone Species. in Biodiversity and Ecosystem Function 237–253 (Springer, Berlin, Heidelberg, 1994). doi:10.1007/978-3-642-58001-7_11

19. Banerjee, S., Schlaeppi, K. & Heijden, M. G. A. van der. Keystone taxa as drivers of microbiome structure and functioning. Nat. Rev. Microbiol. 16, 567–576 (2018).

20. Stewart, F.M. & Levin, B. R. Partitioning of Resources and the Outcome of Interspecific Competition: A Model and Some General Considerations. Am. Nat. 107, 171–198 (1973).

21. Hastings, A. Population Biology: Concepts and Models. (Springer Science & Business Media, 2013).

22. Meers, J. L. Effect of Dilution Rate on the Outcome of Chemostat Mixed Culture Experiments. Microbiology 67, 359–361 (1971).

23. Sommer, U. Phytoplankton competition along a gradient of dilution rates. Oecologia 68, 503–506 (1986).

24. Spijkerman, E. & Coesel, P. F. M. Competition for Phosphorus Among Planktonic Desmid Species in Continuous-Flow Culture1. J. Phycol. 32, 939–948 (1996).

25. Gause, G. F. The Struggle for Existence. (Courier Corporation, 2003).

26. Slobodkin, L. B. Experimental Populations of Hydrida. J. Anim. Ecol. 33, 131–148 (1964).

27. Slobodkin, L. B. Growth and regulation of animal populations. Growth Regul. Anim. Popul. (1980).

28. Friedman, J., Higgins, L.M. & Gore, J. Community structure follows simple assembly rules in microbial microcosms. Nat. Ecol. Evol. 1, 0109 (2017).

29. Celiker, H. & Gore, J. Clustering in community structure across replicate ecosystems following a long-term bacterial evolution experiment. Nat. Commun. 5, 4643 (2014).

30. Vet, S. et al. Bistability in a system of two species interacting through mutualism as well as competition: Chemostat vs. Lotka-Volterra equations. PLOS ONE 13, e0197462 (2018).

31. Kurihara, Y., Shikano, S. & Toda, M. Trade-Off between Interspecific Competitive Ability and Growth Rate in Bacteria. Ecology 71, 645–650 (1990).

32. Luckinbill, L. S. Selection and the r/K Continuum in Experimental Populations of Protozoa. Am. Nat. 113, 427–437 (1979).

33. Andersson, D.I. & Levin, B. R. The biological cost of antibiotic resistance. Curr. Opin. Microbiol. 2, 489–493 (1999).

34. Gross, K. L. Effects of Seed Size and Growth Form on Seedling Establishment of Six Monocarpic Perennial Plants. J. Ecol. 72, 369–387 (1984).

35. Geritz, S. A. H., van der Meijden, E. & Metz, J. A. J. Evolutionary Dynamics of Seed Size and Seedling Competitive Ability. Theor. Popul. Biol. 55, 324–343 (1999).

36. Manhart, M., Adkar, B.V. & Shakhnovich, E. I. Trade-offs between microbial growth phases lead to frequency-dependent and non-transitive selection. Proc R Soc B 285, 20172459 (2018).

37. Venema, K. & van den Abbeele, P. Experimental models of the gut microbiome. Best Pract. Res. Clin. Gastroenterol. 27, 115–126 (2013).

38. Avrani, S., Bolotin, E., Katz, S. & Hershberg, R. Rapid Genetic Adaptation during the First Four Months of Survival under Resource Exhaustion. Mol. Biol. Evol. 34, 1758–1769 (2017).

39. Venturelli, O. S. et al. Deciphering microbial interactions in synthetic human gut microbiome communities. Mol. Syst. Biol. 14, e8157 (2018).

40. Guo, X. & Boedicker, J. Q. The Contribution of High-Order Metabolic Interactions to the Global Activity of a Four-Species Microbial Community. PLOS Comput. Biol. 12, e1005079 (2016).

41. Billick, I. & Case, T. J. Higher Order Interactions in Ecological Communities: What Are They and How Can They be Detected? Ecology 75, 1529–1543 (1994).

42. Bairey, E., Kelsic, E.D. & Kishony, R. High-order species interactions shape ecosystem diversity. Nat. Commun. 7, 12285 (2016).

43. Grilli, J., Barabás, G., Michalska-Smith, M.J. & Allesina, S. Higher-order interactions stabilize dynamics in competitive network models. Nature 548, 210–213 (2017).

44. Mayfield, M.M. & Stouffer, D. B. Higher-order interactions capture unexplained complexity in diverse communities. Nat. Ecol. Evol. 1, 0062 (2017).

45. Goldford, J. E. et al. Emergent simplicity in microbial community assembly. Science 361, 469–474 (2018).

46. Niehaus, L. et al. Microbial coexistence through chemical-mediated interactions. bioRxiv 358481 (2018). doi:10.1101/358481

47. Fox, J. W. The intermediate disturbance hypothesis should be abandoned. Trends Ecol. Evol. 28, 86–92 (2013).

48. Chesson, P. & Huntly, N. The Roles of Harsh and Fluctuating Conditions in the Dynamics of Ecological Communities. Am. Nat. 150, 519–553 (1997).

49. Hsu, S.-B. & Zhao, X.-Q. A Lotka–Volterra competition model with seasonal succession. J. Math. Biol. 64, 109–130 (2012).

50. Tresse, O., Jouenne, T. & Junter, G.-A. The role of oxygen limitation in the resistance of agar-entrapped, sessile-like Escherichia coli to aminoglycoside and β-lactam antibiotics. J. Antimicrob. Chemother. 36, 521–526 (1995).

51. Ratkowsky, D. A., Olley, J., McMeekin, T.A. & Ball, A. Relationship between temperature and growth rate of bacterial cultures. J. Bacteriol. 149, 1–5 (1982).

